# A ratiometric dual-color luciferase reporter for fast characterization of transcriptional regulatory elements

**DOI:** 10.1101/2021.05.26.443018

**Authors:** E. González-Grandío, G. S. Demirer, W. Ma, S.M. Brady, M.P. Landry

## Abstract

Plant synthetic biology requires precise characterization of genetic elements to construct complex genetic circuits that can improve plant traits or confer them new characteristics. Transcriptional reporter assays are essential to quantify the effect of gene expression regulator elements. Therefore, transcriptional reporter systems are a key tool in understanding control of gene expression in biology.

In this work we construct and characterize a dual-color luciferase ratiometric reporter system that possesses several advantages over currently used reporters. It is ratiometric, reducing variability and increasing consistency between experiments; it is fast, as both reporters can be measured at the same time in a single reaction, and it is cheaper to perform than current dual-luciferase reporter assays. We have validated our system quantifying the transcriptional capability of a panel of promoters and terminators commonly used in synthetic biology with a broad range of expression magnitudes, and in a biologically relevant system, nitrate response.

## Introduction

Agronomy faces significant global challenges and addressing these will require the adoption of genetic manipulation technologies used in synthetic biology. Synthetic biology tools can aid in both understanding endogenous transcriptional regulation and designing genetic circuits to confer plants novel traits. Complex DNA constructs can be assembled from different genetic elements with modular cloning toolkits (Vazquez-Vilar *et al.*, 2017) together with the Phytobrick standard that systematizes sequence requirements for cloning (Patron *et al.*, 2015). In plants, the most commonly employed reporters to measure transcriptional activity are fluorescent proteins, beta-glucuronidase, and luminescent proteins (De Ruijter *et al.*, 2003). Variability commonly found between samples and experiments can be reduced using ratiometric assays with a second invariable reporter as an internal reference (Vazquez-Vilar *et al.*, 2017). However, activity of these dual reporters is measured by different methods, or in sequential reactions with distinct kinetics (Sarrion-Perdigones *et al.*, 2019). Additionally, these assays often require sample homogenization and protein extraction, and are thus unpractical for high-throughput assays.

Here, we developed a ratiometric dual-color luciferase reporter assay to quantify transcriptional activity of genetic elements in plants. This GREAT (<u>G</u>reen/<u>Re</u>d luciferase r<u>at</u>iometric) reporter system is based on two luciferases that process the same substrate and emit light at different wavelengths. This system is i) ratiometric, with both reporters measured by the same method and at the same time, ii) has a high dynamic range output that allows precise comparison of a wide range of expression magnitudes, iii) requires minimal sample preparation time and, iv) is cheaper than currently used dual luciferase methods.

## Results and Discussion

We surveyed a panel of luciferases that emit green or red luminescence (Sarrion-Perdigones *et al.*, 2019) by transiently expressing them in *Nicotiana benthamiana* leaves through agroinfiltration as in (Vazquez-Vilar *et al.*, 2017). We extracted total protein from leaf tissues and measured luciferase emission spectra and stability three days post-infiltration using the Promega Steady-Glo Assay, in a Tecan M-100PRO plate reader. We selected E-Luc (Nakajima *et al.*, 2010) and Red-F (Branchini *et al.*, 2006) as green- and red-emitting luciferases, respectively, because this pair of luciferases exhibited the most distant emission peaks (Figure 1A, 541 nm and 608 nm, respectively). Additionally, both luciferases showed very similar and stable reaction kinetics that allow reliable relative measurements starting 10 minutes after extraction and substrate mixing, with stable luminescence for over 2 hours (Figure 1B). These properties allow characterization of many samples in a standard luminometer without the need for expensive injectors required for time-sensitive coelenterazine-using luciferases (e.g., Renilla luciferase). Moreover, we were able to measure stable luminescence directly from leaf discs (Figure 1B). This method greatly reduces sample processing time and labor, and thus, is amenable for high-throughput measurements.

**Figure 1.**
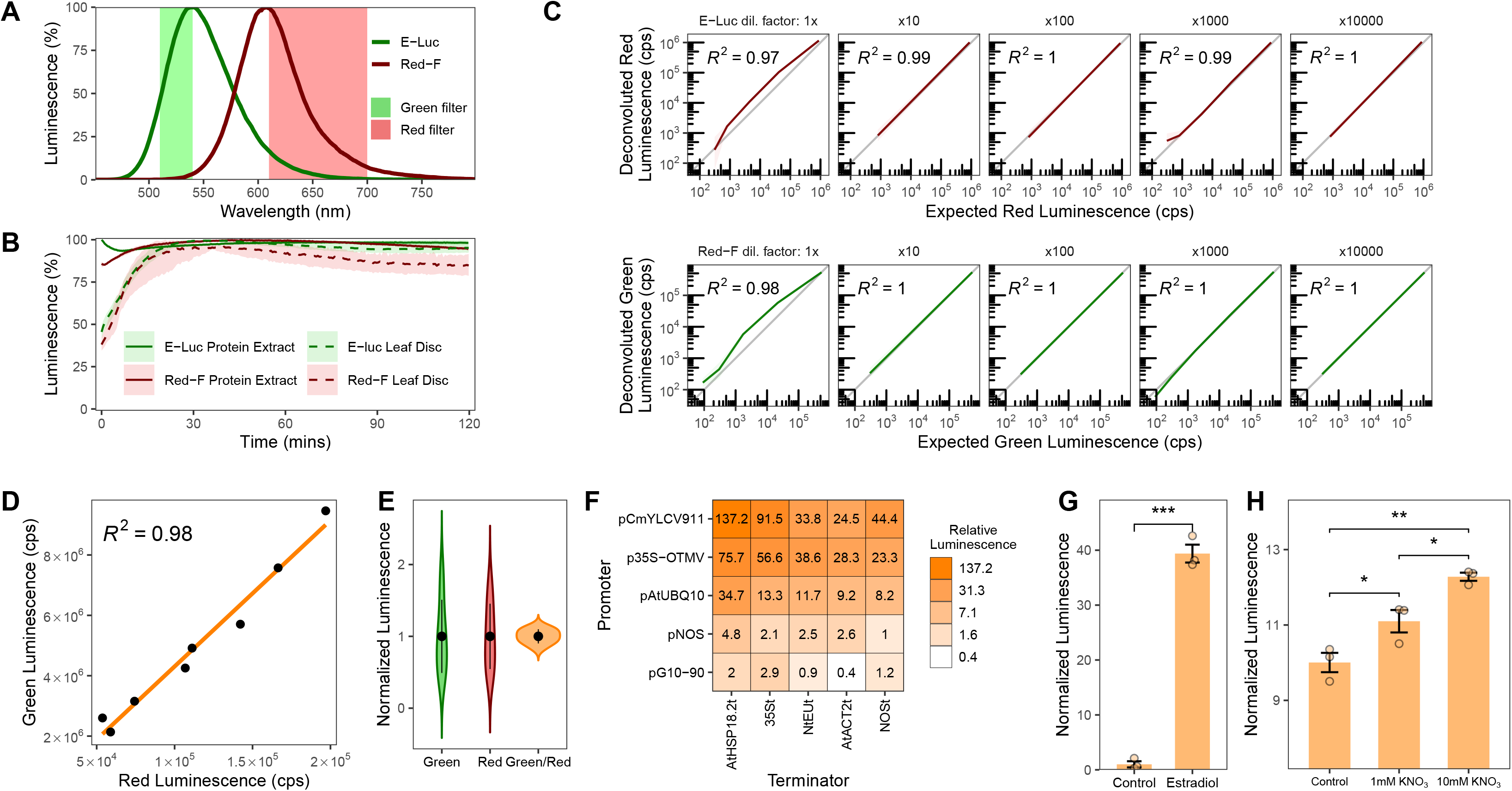
Characterization of a ratiometric dual-luciferase reporter system. A) Emission spectra of E-Luc and Red-F luciferases. Wavelengths transmitted by green- and red-light filters are shown. B) Luminescence stability from protein extract or leaf discs. C) Signal deconvolution of Red-F and E-Luc. Deconvoluted signal is compared to expected red or green luminescence in counts per second (cps). D) Correlation of deconvoluted green and red luminescence from plants agroinfiltrated with pGREAT2. The orange line represents linear regression. E) Dispersion of luminescence in samples from panel D. Green and red luminescence are normalized to the average luminescence. The black dot represents the average and vertical lines the standard deviation. F) Green/red luminescence from plants agroinfiltrated with pGREAT1-25 plasmids. Green/red luminescence is relative to pGREAT1 (pNOS:E-Luc:NOSt). G) Green/red luminescence of an agroinfiltrated plasmid in which E-luc is under control of an estradiol inducible promoter (pGREAT26). Leaves were infiltrated with 20 μM β-estradiol. H) Green/red luminescence of Arabidopsis root protoplasts transformed with a plasmid in which E-luc is under control of the *Nitrate-Regulated Promoter* (pGREAT27). Protoplasts were incubated with increasing nitrate concentrations. In panels B and C, the line represents average luminescence, the faded band represents a 95% confidence interval. Error bars in G and H represent the standard error of the mean. T-test statistical significance in G and H is denoted as follows: *, p-val < 0.05; **, p-val < 0.01; ***, p-val < 0.001. n = 3 for each treatment/condition except in panel D and E, where n = 8 and panel C, where n = 3-6.

As there is partial signal overlap between E-Luc and Red-F, we used a previously-described simple deconvolution method that allows precise adscription of luminescence signal to each luciferase (Promega Technical Manual TM062). Calibration constants are calculated measuring green-, red-filtered, and total luminescence (Figure 1A). These constants only need to be calculated once if the measurement conditions that affect luminescence are kept fixed in subsequent experiments (e.g., temperature, pH). We tested signal deconvolution for each luciferase by measuring luminescence of a serial dilution of one luciferase to which we added different amounts of the other luciferase. We then compared the deconvoluted signal to expected red or green luminescence measured from extracts with only one of the luciferases. We found that when luminescence from the second luciferase is very high compared to the first one, deconvoluted signal of the first luciferase can be overestimated (Figure 1C, leftmost panels). When the amount of the second luciferase was reduced, this overestimation was eliminated and the deconvoluted luminescence perfectly matched the expected luminescence value (Figure 1C). Based on these results, we selected a Red-F transcriptional unit that produced moderately low luminescence (pNOS:Red-F:NOSt) as the normalizer reporter for our ratiometric system. We recommend conducting a similar assay to the one described here and including the deconvolution deviation in final calculations when the expected measurements will vary over a wide range of magnitudes.

We then constructed a ratiometric vector including both luciferases to quantify how ratiometric normalization reduces variability when measuring transcriptional activity (pGREAT2). We agroinfiltrated several *Nicotiana benthamiana* leaves and, as expected, we observed a great degree of variability when measuring green luminescence in different leaves, quantified by a coefficient of variability (CV) of 50.93%. This variability was also proportional to red luminescence (CV of 45.44%), suggesting that each leaf exhibited diverse degrees of transcription/translation that affected both luciferases equally (Figure 1D). When green luminescence was normalized by red luminescence, variability was greatly reduced to a CV of 9.85% (Figure 1E).

We then used the GREAT reporter to systematically measure transcriptional activity of a set of five promoters and five terminators commonly used in plant synthetic biology. We specifically chose genetic elements with different activities to test our system over a wide range of expression levels. We Phytobrick-adapted these sequences, and built E-Luc transcriptional units with all possible combinations of promoters and terminators together with pNOS:Red-F:NOSt for ratiometric normalization, using the modular GoldenBraid system (Vazquez-Vilar *et al.*, 2017) (pGREAT1-25). We agroinfiltrated these vectors into *Nicotiana* leaves and measured green/red luminescence of each plasmid, using the pGREAT1 (pNOS:E-Luc:NOSt) vector as an internal normalizer, as this practice has been shown to reduce inter-experiment technical variability (Vazquez-Vilar *et al.*, 2017). We found that our system enabled expression quantification from 0.4 to 137.2 relative luminescence units, confirming that our system can be used to reliably quantify transcriptional activity over at least 4 orders of magnitude (Figure 1F). The aforementioned characteristics of the GREAT reporter system are ideal for these types of screens by reducing the time and cost of the assay. Additionally, we used our system to measure the responsiveness of a Phytobrick-adapted β-estradiol inducible promoter (pGREAT26) (Zuo *et al.*, 2000). We were able to measure a 40-fold green/red luminescence induction 24h after treatment with 20 μM β-estradiol (Figure 1G).

We also tested if the GREAT system could be used to measure biologically-relevant transcriptional responses. We cloned the Arabidopsis *Nitrate-Regulated Promoter* (Wang *et al.*, 2010) to control E-Luc expression (pGREAT27). We transformed Arabidopsis root protoplasts (Birnbaum *et al.*, 2005; Yoo *et al.*, 2007) and treated them with increasing nitrate concentrations for 16h. We observed a significant induction of green/red luminescence proportional to the nitrate concentration, validating our system (Figure 1H).

In summary, we have developed a system that possesses several advantages over other commonly-used methods to measure transcriptional activity, and we have provided evidence of its potential applications. We have deposited the standardized Phytobricks and pGREAT1-27 plasmids generated herein in the Addgene repository, including detailed sequence maps (Addgene #170873-170915). We expect that the system described in this work will help advance our understanding of transcriptional regulation and accelerate plant synthetic biology.

## Author Contributions

E.G. and M.P.L. conceived the project and wrote the manuscript. E.G., G.S.D., S.M.B. and M.P.L. designed the experiments. E.G. performed the experiments and data analysis. E.G. and W.M. prepared the plasmids. G.S.D. performed the nitrate experiments. All authors have edited and commented on the manuscript and have given their approval of the final version.

## Acknowledgements

M.P.L acknowledges support of a USDA BBT EAGER award (2019-67013-29104) and a USDA NIFA Award (2018-67021-27964). G.S.D is supported by the Resnick Sustainability Institute Postdoctoral Fellowship, and G.S.D and S.M.B. acknowledge support of a USDA BBT-EAGER award (2019-67013-29012), and partial support by an HHMI Faculty Scholar fellowship. pRedF was a gift from Koen Venken (Addgene #118057).

